# Estimating statistical significance of local protein profile-profile alignments

**DOI:** 10.1101/484485

**Authors:** Mindaugas Margelevičius

## Abstract

Alignment of sequence families described by profiles provides a sensitive means for establishing homology between proteins and is important in protein evolutionary, structural, and functional studies. In the context of a steadily growing amount of sequence data, estimating the statistical significance of alignments, including profile-profile alignments, plays a key role in alignment-based homology search algorithms. Still, it is an open question as to what and whether one type of distribution governs profile-profile alignment score, especially when profile-profile substitution scores involve such terms as secondary structure predictions. This study presents a methodology for estimating the statistical significance of this type of alignments. The methodology rests on a new algorithm developed for generating random profiles such that their alignment scores are distributed similarly to those obtained for real unrelated profiles. We show that improvements in statistical accuracy and sensitivity and high-quality alignment rate result from statistically characterizing alignments by establishing the dependence of statistical parameters on various measures associated with both individual and pairwise profile characteristics. Implemented in the COMER software, the proposed methodology yielded an increase of up to 34.2% in the number of true positives and up to 61.8% in the number of high-quality alignments with respect to the previous version of the COMER method. A new version (v1.5.1) of the COMER software is available at https://sourceforge.net/projects/comer. The COMER software is also available on Github at https://github.com/minmarg/comer and as a Docker image (https://hub.docker.com/r/minmar/comer).

## 1 Introduction

Establishing homology between proteins is essential in evolutionary and highly important in structural and functional studies of proteins and their complexes. Alignment, in general, represents the most fundamental way to deduce homology directly or indirectly. Sequence alignment has proved indispensable in annotating uncharacterized proteins and paved the way for alignment of sequence families described by profiles, constituting the basis for inferring protein structure and function by homology.

In the presence of a large and steadily growing amount of sequence data, estimating the statistical significance of alignments plays a prominent role in alignment-based homology search algorithms (Karlin, 2005). For an alignment with a particular similarity score, it provides a probability or related quantity indicating how likely a chance alignment with the same or greater score is to be observed.

The limiting distribution of the ungapped local alignment score *S* (nonlattice) for large sequence lengths *m* and *n* has been proved (Karlin *et al.*, 1990; Karlin and Altschul, 1990; Karlin and Brendel, 1992; Dembo *et al.*, 1994) to be the type 1 (Gumbel-type) extreme value distribution (EVD) (Kotz and Nadarajah, 2000)

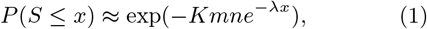

where *λ* and *K* (*K* = *e*^*λμ*^/[*mn*]; *μ*, the location parameter under the standard parametrization) are parameters calculable from the score matrix used to align sequences and satisfying the conditions of the negative expected score and existence of at least one positive score.

Ample empirical evidence (e.g., Mott, 1992; Altschul and Gish, 1996; Pearson, 1998; Waterman and Vingron, 1994b,a) has suggested that the statistical theory for ungapped alignments applies to gapped alignments, provided gap penalties, or costs, lead to alignment scores in the (local) region of logarithmic growth (Arratia and Waterman, 1994).

Importantly, factors, such as the length of sequences being compared (Altschul and Gish, 1996; Spang and Vingron, 1998; Altschul *et al.*, 2001) and their compositional similarity (Mott, 1992, 2000), affect the distribution of alignment scores. While solutions to take them into account exist for sequence and profile-to-sequence alignments (Mott, 1992; Yu *et al.*, 2006) (Supplementary Section S1), there is no established procedure for addressing them in profile-profile alignment (Sadreyev and Grishin, 2003; Poleksic, 2009; Margelevičius and Venclovas, 2010; Sadreyev and Grishin, 2008).

This study aims at characterizing the distribution of profile-profile alignment scores to improve statistical accuracy and remote homology detection. Our focus is local alignment scores. Hence, we assume that the expected profile-profile alignment score per aligned pair is negative and the probability for a positive score is positive. These assumptions are satisfied in part when the score for a pair of positions of two profiles is (implicitly) log-odds (Karlin, 2005; Meng *et al.*, 2011). Importantly, the score preserves the form of log-odds when it linearly combines different components (e.g., the similarity of two amino acid frequency vectors with that of secondary structures) as long as each component itself represents a log-odds score. Such a construction of composite scores is typical, and, given appropriate gap penalties, we can expect that the assumptions will hold for most profile-profile alignment methods.

Here, we use the COMER method (Margelevičius, 2016) as a means for producing profile-profile alignment scores the algorithms developed in this work employ. The COMER profile at each position encapsulates the transition probabilities of moving to and from the states of insertion and deletion, which for a pair of profiles transform to position-specific gap penalties. Comparing two COMER profiles also includes scoring predicted secondary structure (SS) and sequence contexts (Margelevičius, 2018). These properties make the characterization of the distribution of alignment scores a challenging task.

Moreover, the distribution of profile-profile alignment scores strongly depends on the extent of (dis)similarity between unrelated profiles, or the null model of random sequence families. Real sequences do not conform to the canonical (and simplest) model, in which the amino acids in the sequence are iid, and exhibit a more complex structure with short- and long-range amino acid correlations (Mott, 2000; Yu and Hwa, 2001; Metzler, 2006; Eddy, 2008; Meng *et al.*, 2011). It has been proved that in the limit of infinitely long sequences, ungapped alignment scores for random Markov-dependent sequences are still distributed according to the EVD (Karlin and Dembo, 1992). Gapped alignments of correlated random sequences accounted for by a null model have been shown numerically to correspond to an EVD too (Messer *et al.*, 2007). Notably, the values of the statistical parameters differed substantially from those obtained for iid sequences.

In this work, the null model of protein sequence family does not explicitly incorporate correlations between families and leaves the profile-profile scoring function unchanged. Instead, we randomly generate profiles with properties of real unrelated profiles and take into account the correlations between them by establishing the dependence of statistical parameters on the compositional similarity between the profiles. The null model affects the probability of a score of a pair of profile positions, yet the change in composition does not alter the type of alignment score distribution but values of statistical parameters. The approach proposed here predicts differences in these values.

Overall, this work purposes a procedure for estimating the statistical significance of profile-profile alignments, regarding the effects and factors that influence alignment score distribution, such as edge effects, compositional similarity, and profile attributes (length and the effective number of observations). The procedure builds on an algorithm proposed for generating profiles that resemble real sequence families. The results of the implemented procedure give rise to an analysis of its impact on sensitivity and alignment quality, which we also provide in this paper.

## 2 Methods

The concept of estimating the statistical significance of profile-profile alignments, as proposed here, is illustrated in Figure 1.

The significance of the alignment between a profile of length *l*_1_ and effective number of observations (ENO) *n*_1_ and another profile of length *l*_2_ and ENO *n*_2_ follows from combining the significance of the alignment score and that of the number of positive substitution scores in the alignment. The distribution of alignment scores depends on both the lengths of profiles being compared and their ENOs (Sadreyev and Grishin, 2008) (Supplementary Section S2.1 defines ENO, which represents the median number of residues per profile position). The composition of profiles is another factor that affects the distribution of alignment scores, and it is a measure independent of profile ENO. Therefore, we incorporate a measure of compositional similarity calculated between profiles into a statistical model for the distribution of alignment scores. The significance of the alignment score of profiles 1 and 2 is then calculated using statistics obtained from aligning unrelated profiles of the same length and ENO and having the same mutual compositional similarity.

The parameters of alignment score distributions are estimated by simulation. We develop algorithms to generate randomly profiles that express properties of, and their alignment scores distribute similarly to alignment scores of, real unrelated profiles. Dividing random profiles according to (1) their length and ENO and (2) length and mutual compositional similarity provides two categories of profiles. Two categories of distributions of alignment scores, illustrated in Figure 1, result from aligning the profiles in each category. We develop conditional mean estimators to combine the statistics of these distributions, which captures the dependence of the distribution of alignment scores on profile length, ENO, and the compositional similarity between profiles. Finally, we derive a statistic based on the number of positive profile-profile substitution scores and combine its statistical significance and the significance of the alignment score to obtain the final estimate.

**Figure 1.**
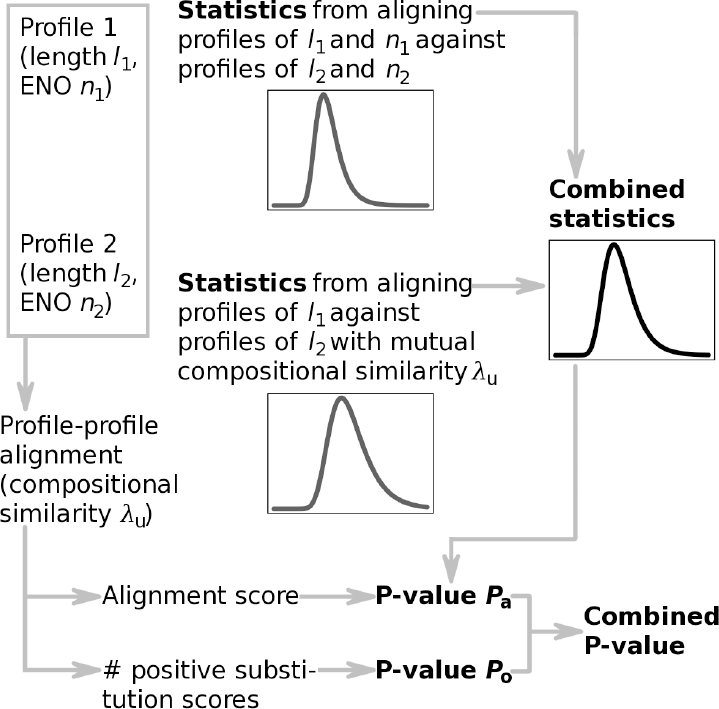
Concept of estimating the statistical significance of profile-profile alignments.

These steps are described in detail below. More details can be found in Supplementary Section S4.

### 2.1 Compositional similarity

The statistical parameter *λ*_u_ ≡ *λ* of the limiting distribution of sequence alignment scores (1) has several related meanings (Karlin and Altschul, 1990; Altschul *et al.*, 1997; Schäffer *et al.*, 2001). *λ*_u_ can also be regarded as a measure of compositional similarity (Yu *et al.*, 2006). The value of *λ*_u_ found as the positive solution to

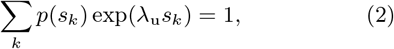

where {*s*_*k*_}_*k*_ represent different values of scores in the substitution matrix and *p*(*s*_*k*_) is the probability of *s*_*k*_, will decrease as the number of positive substitution scores increases (*p*(*s*_*k*_) increases for all *k*: *s*_*k*_ > 0). Therefore, low values of *λ*_u_ may indicate an increased probability for a high-scoring alignment to occur by chance due to compositionally biased regions in two sequences.

The parameter *λ*_u_ calculated for a pair of profiles (Sadreyev and Grishin, 2003; Margelevičius and Venclovas, 2010) has the same dependence on composition. We take into account compositional similarity between profiles by specifying the dependence of the distribution of profile-profile alignment scores on it.

### 2.2 Alignment scores of real unrelated profiles

We analyze the distribution of alignment scores of real unrelated profiles (constructed from multiple sequence alignments, MSAs, of real sequences) for two reasons. One is to determine the type of distribution that describes them. The other reason is that these data provide a reference point for generating random profiles whose alignment scores would follow the same type of distribution and be similarly distributed.

We found that alignment scores of real unrelated profiles follow an EVD and 85% of goodness-of-fit tests do not reject the null hypothesis. Supplementary Sections S2.3 and S3.1, respectively, describe how real unrelated profiles were obtained and detail the comparison of profiles of different length, ENO, and compositional similarity.

### 2.3 Profile simulation

It is impractical to use real unrelated profiles to produce sufficient data for any values of profile length, ENO, and compositional similarity. Instead, random profiles are generated.

It has been shown that using a null model to generate realistic random profiles improves remote homology detection (Sadreyev and Grishin, 2008). However, using the earlier procedure for generating random profiles may lead to highly correlated profiles when the comparison of generated profiles includes the scoring of predicted SSs (Supplementary Section S3.2). Alignment scores of random profiles may not then represent a distribution obtained by aligning unrelated profiles. Therefore, the aim is to develop an algorithm for generating random profiles that exhibit properties of real profiles, and the degree of correlation between the random profiles can be controlled.

We achieve that by using a modest number of “seed” profiles constructed for a diverse set of real sequences. Random profiles then result from adding noise to profiles/MSAs generated using these seed profiles as a model. Seeds provide properties of real profiles, whereas noise and randomization represent a means of controlling correlation between random profiles.

An important feature of the algorithm that we propose (Algorithm 1) is that random profiles result from concatenating fixed-length fragments sampled randomly from a set of generated MSAs regardless of SS predictions the fragments entail. Supplementary Section S4.1 provides additional details, and Supplementary Algorithm S2 defines how MSAs with added noise are produced using a profile model in step 2 of Algorithm 1.

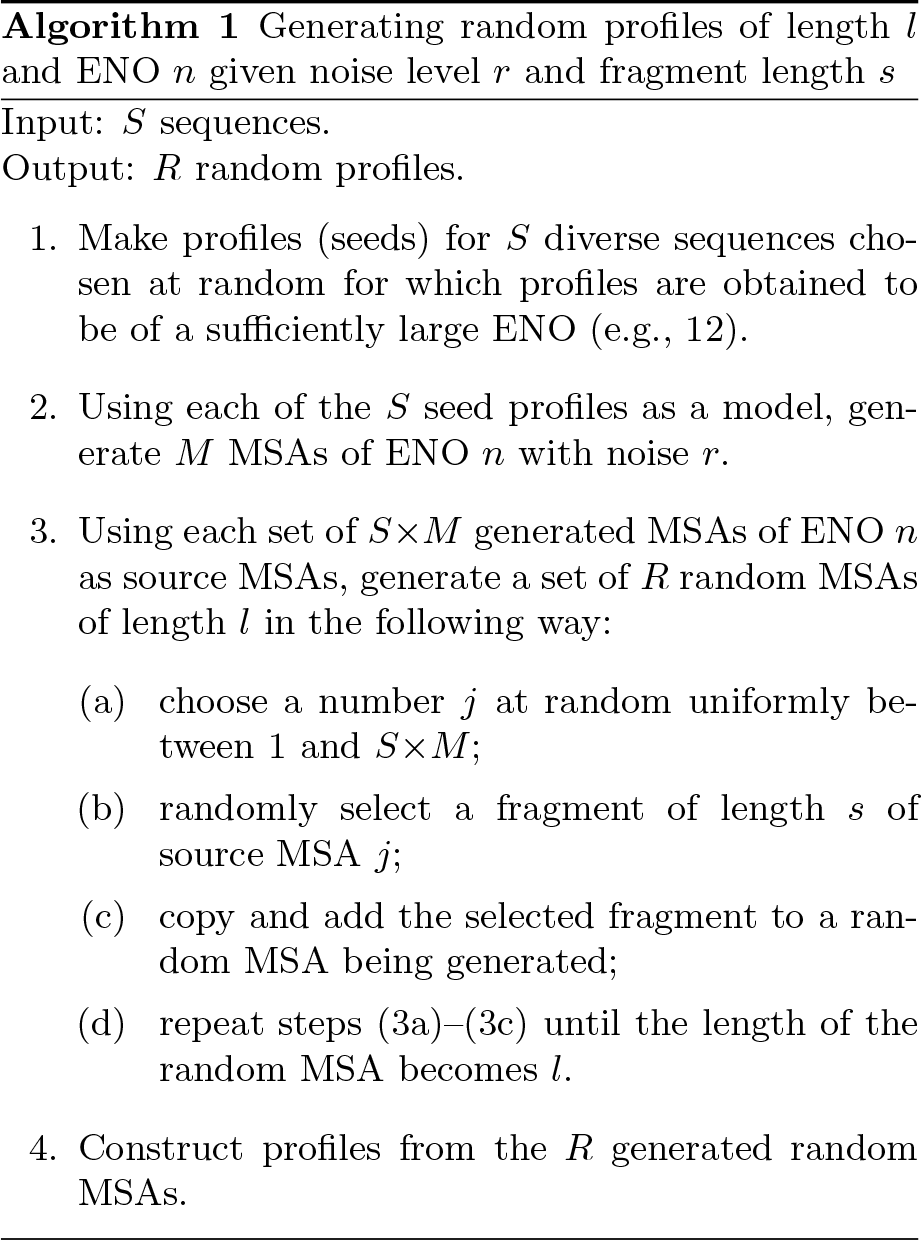

Algorithm 1 shows that the fragment length *s* and the level *r* of noise added to *S*×*M* generated MSAs are the parameters that determine the degree of similarity among resulting random profiles. Too large *s* and/or too small *r* lead to highly correlated random profiles, while too small *s* or too large *r* result in divergent profiles and useless alignment statistics.

### 2.4 Optimizing *s* and *r*

Based on the results reported in Section 2.2, we find the optimal *s* and *r* using two criteria: (1) the goodness of fit of the EVD to the empirical distribution of alignment scores and (2) the distance between the distribution function obtained for real unrelated profiles and the distribution function obtained from simulations. In this way, profiles generated using the optimal *s* and *r* possess features characteristic to real profiles and correlations between them do not dominate.

We calculate the supremum class upper tail Anderson-Darling statistic *AD*_up_ (Chernobai *et al.*, 2015) (Supplementary Section S5.1) for testing the goodness of fit of the EVD to the data (the first criterion). We normalize it, 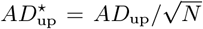, by the square root of the number of alignment scores, 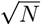, to make it independent of sample size. The distance bet-ween two empirical distribution functions (the second criterion) is measured by the two-sample Kolmogorov-Smirnov statistic *D*.

The whole procedure for finding the optimal *s* and *r* can be described as follows. 1. Choose values for *s* and *r*. 2. Using Algorithm 1, generate random profiles of all ENOs and lengths considered. 3. Align simulated profiles. 4. Fit the EVD in the right tail of alignment score distributions. 5. Calculate the 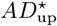 statistic for each alignment score distribution. 6. Train models for predicting the statistical parameters for profiles with given attributes (ENO and length) and compositional similarity. 7. Predict the statistical parameters and calculate the *p*-value of each alignment of real unrelated profiles. 8. For each combination of pair values of profile ENO and length, calculate the distance *D* between the empirical distribution function obtained for real unrelated profiles and the distribution function obtained in step 7. 9. Repeat steps 1–8 until the optimal *s* and *r* with respect to 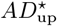 and *D* have been found.

We discuss prediction of statistical parameters (step 7) below. Here we note that predictions are used to incorporate compositional similarity into the statistical model of profile-profile alignments.

The results (Figure 2A) obtained using *S* = 1012 seed profiles (*M* = 1) suggest that considering a small number of different values suffices to find the optimal *s* and *r*: Large *s* increases correlation between profiles, whereas large *r* makes them unalignable. Figure 2A shows that the best balance between the two criteria is achieved for *s* = 9 and *r* = 0.03.

Sections S6.1 and S6.2 present additional results from simulation experiments. They also show that seeding from three different profiles representing different SCOPe (Fox *et al.*, 2013) classes leads to similar results (Figure 2B). Using one seed facilitates profile simulation.

### 2.5 Conditional mean estimators

Consider two profiles, one of length *l*_1_ and ENO *n*_1_ and another one of length *l*_2_ and ENO *n*_2_. The profiles share compositional similarly *λ*_u_. Then, the significance of their alignment score is determined from combining statistics obtained (1) from aligning simulated profiles of length *l*_1_ and ENO *n*_1_ against profiles of length *l*_2_ and ENO *n*_2_ and (2) from aligning profiles of length *l*_1_ against profiles of length *l*_2_ with mutual compositional similarly *λ*_u_, as shown in Figure 1. Here we define this statistical combination.

**Figure 2.**
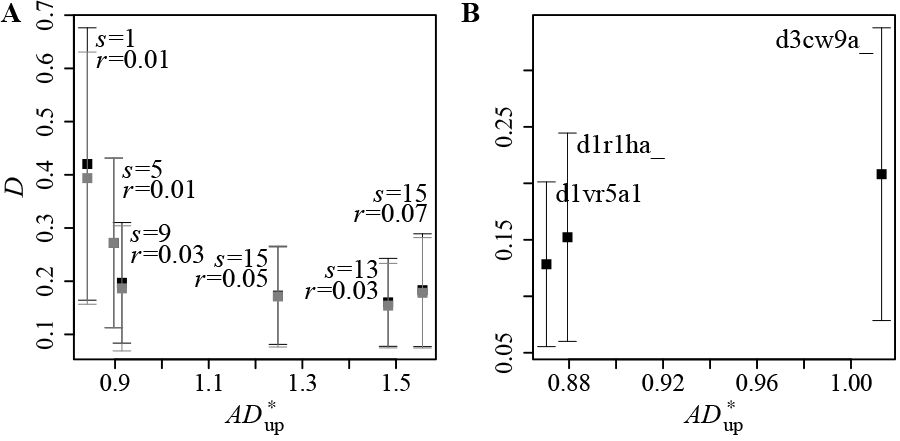
Distance between distributions against goodness of fit for different values of *s* and *r*. Each point represents the average over all distributions obtained for different pair values of profile attributes. Vertical bars represent one standard deviation. Gray and black colors show the results obtained with and without considering the compositional similarity between profiles, respectively. (A): Results obtained using *S* = 1012 seed profiles. (B): Results for three different seed profiles (*S* = 1) used to generate *M* = 1000 source MSAs with *s* = 9 and *r* = 0.03. In this case, the results obtained with and without considering compositional similarity coincide.

Let A and B index two distributions. The first is obtained for profiles described by the set of attributes {*n*_1_, *l*_1_; *n*_2_, *l*_2_} (i.e., profiles of length *l*_1_ and ENO *n*_1_ have been aligned against profiles of length *l*_2_ and ENO *n*_2_) and the other for profiles characterized by the set of attributes {*λ*_u_; *l*_1_; *l*_2_}. Assume that distributions A and B belong to the family of EVDs. Let 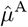 and 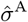 respectively denote the estimates of the location and scale parameters of the EVD corresponding to distribution A. Similarly, let 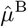 and 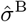 be the estimates of the statistical parameters of distribution B. Then, the conditional mean estimator of the location parameter given two sets of parameters specifying its distribution in settings A and B is

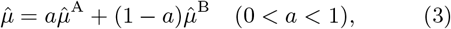

and the corresponding conditional mean estimator of the scale parameter is

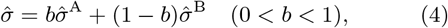

where *a* and *b* are parameters that depend on the parameters specifying the distributions of the location and scale parameters, respectively, in settings A and B (see Supplementary Appendix A for details and a proof).

### 2.6 Prediction of statistical parameters

In simulations (Section 2.4), random profiles are generated of predefined lengths *l* ∈ **L** and ENOs *n* ∈ **N** and discretized values of mutual compositional similarity *λ*_u_ rounded to the nearest multiple of 0.1. The sets **L** = {50, 100, 200, 400, 600, 800} and **N** = {2, 4, 6, …, 14} represent these predefined values.

**Figure 3.**
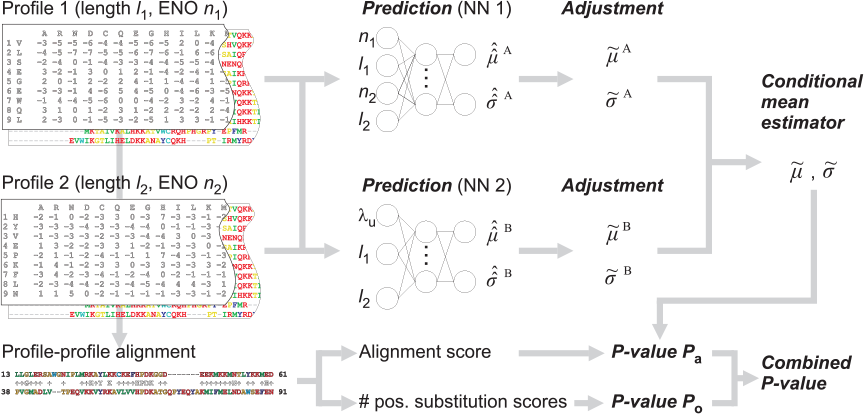
Complete procedure of estimating the statistical significance of profile-profile alignments.

Statistical significance for profiles of any length *l*, ENO *n*, and *λ*_u_ has to be estimated based on the statistics obtained from the observed distributions. As the statistics depend non-linearly on *l*, *n*, and *λ*_u_ (Supplementary Section S6), we use low-complexity artificial neural network (NN) models to predict statistical parameters. The models are trained on the estimates obtained (1) from aligning simulated profiles of length *l*_1_ ∈ **L** and ENO *n*_1_ ∈ **N** against profiles of length *l*_2_ ∈ **L** and ENO *n*_2_ ∈ **N** (*n*_2_ < *n*_1_) and (2) from aligning profiles of length *l*_1_ ∈ **L** against profiles of length *l*_2_ ∈ **L** with *λ*_u_ discretized. We denote the predictions of the location parameter in these two settings by 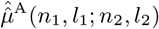 and 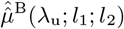, respectively. The notation for the scale parameter is similar. (See also Supplementary Section S4.4.)

### 2.7 Adjustment of predictions

Statistical accuracy does not necessarily correlate with increased sensitivity (Yu *et al.*, 2006; Sadreyev and Grishin, 2008) and high-quality alignment rate. Therefore, to simultaneously improve all these qualities, we introduce simple adjustments to the predicted location 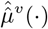 and scale parameters 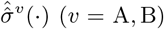, as shown in Figure 3.

Observing a high correlation between the scale and the location parameters (Supplementary Section S6.3), we adjust the scale parameters as follows:

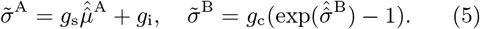

The first equation eliminates the need to predict 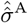 The second equation, although expressed non-linearly in 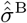, employs one coefficient whose optimal value implies almost the same result as in the case of expressing the scale parameter linearly in 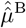.

We adjust the location parameters similarly:

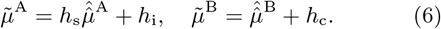

Note that the adjustments 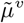 and 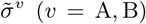 retain the dependence of the statistical parameters on the profile length and ENO and compositional similarity. They only scale and shift predictions made by the trained models.

Conditional mean estimators (3) and (4) are used to combine 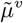 and 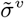. The parameters *a* and *b* and the coefficients *W* = {*g*_*s*_, *g*_*i*_, *g*_*c*_, *h*_*s*_, *h*_*i*_, *h*_*c*_} in (5) and (6), which we refer to as the adjustment parameters, are optimized with respect to statistical accuracy and alignment quality and sensitivity (Supplementary Section S6.5). Here we note that the optimal balance is achieved with the compositional similarity between profiles having a large impact (*a* = *b* = 0.35) on conditional mean estimates.

### 2.8 Number of positive substitution scores

The number of positive profile-profile substitution scores in the alignment provides additional information to the alignment score. For example, the same alignment score can be the result of many weakly positive substitution scores or a few high scores. Therefore, it can be a useful indicator for estimating the significance of profile-profile alignments.

The distribution of the number of positive substitution scores can be accurately approximated by the negative binomial distribution (NBD) (Supplementary Section S6.4). However, aiming to reduce the overall complexity of the calculation of statistical significance, we propose a derived statistic *ω*_n_. It depends on the lengths of profiles being compared and its value (consequently, its significance) decreases as the alignment search space (the product of the profile lengths) increases:

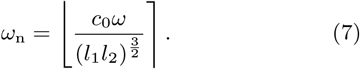

Here ⌊·⌉ denotes rounding to the nearest integer, *l*_1_ and *l*_2_ are the lengths of two profiles compared, *ω* is the number of positive substitution scores in the alignment, and *c*_0_ = 10^5^ is the constant that is equal to the value of the denominator when *l*_1_ and *l*_2_ are slightly less than 50. Hence, the *ω*_n_ statistic corresponds to the number of positive substitution scores when the search space approximately equals that of two profiles of length 50. The number of positive substitution scores becomes less informative as the search space increases, and, consequently, the value of the *ω*_n_ statistic decreases. The distribution of *ω*_n_ is approximately negative binomial (Supplementary Section S6.4).

The complete procedure for statistical significance estimation is shown in Figure 3. *p*-values of alignment score and *ω*_n_, *P*_a_ and *P*_o_, are combined using the empirical Brown’s method (Poole *et al.*, 2016) (Supplementary Section S4.5).

### 2.9 *E*-value and its correction

The expected number of local alignments with a score greater than or equal to *x*, *E* = *Kl*_1_*l*_*N*_*e*^−*λx*^, increases as the size of the search space increases (Karlin and Altschul, 1990; Dembo *et al.*, 1994; Waterman and Vingron, 1994b; Spang and Vingron, 2001). (*E*-value and *p*-value *P* are related by the equation *P* = 1−exp(−*E*).) Let *E*_*N*_ be the *E*-value of an alignment with score *x* obtained from searching a database of size *l*_*N*_. Let also *E*_*N′*_ be the *E*-value of an alignment with the same score *x* (and profile composition) obtained by searching a database of size *l*_*N′*_ with the same query. Then, the relationship between *E*_*N*_ and *E*_*N′*_ is (Yu *et al.*, 2006)

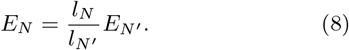

We use this relationship when compensating for a limited number of randomly generated profiles used in simulations. We refer to the results obtained by calculating the corrected *E*-value as the alternative model.

## 3 Results

This section presents results from the application of the proposed procedure (Figure 3) for estimating the statistical significance of profile-profile alignments. The NN models predicting statistical parameters were trained on the estimates obtained for profiles generated by Algorithm 1 with a noise level *r* = 0.03 and a fragment length *s* = 9, as determined in Section 2.4.

For comparison, we have implemented and present results from the application of the method for estimating statistical significance proposed by Sadreyev and Grishin (2008) (Supplementary Section S5.3.4). We refer to the COMER version implementing this method as S&G’08 in the text. We also show the results of the application of the COMPASS (Sadreyev and Grishin, 2003) (v3.1) profile-profile alignment method, where the approach (Sadreyev and Grishin, 2008) applies to the scoring function for which it was originally developed.

### 3.1 Statistical accuracy

We used COMER (or another tool) to search and align the profiles constructed for 5000 simulated Pfam (Finn *et al.*, 2016) (v30.0) MSAs against the database of simulated profiles representing 4931 SCOPe (v2.03) domains. The profiles were randomized using Algorithm 1 to preserve length, ENO, and composition inherent in each of the Pfam MSAs and profiles constructed for the SCOPe domains (see Supplementary Section S5.2). Then, we calculated the fraction of queries with a *p*-value reported by COMER (or another tool) for their best match less than the specified *p*-value.

**Figure 4.**
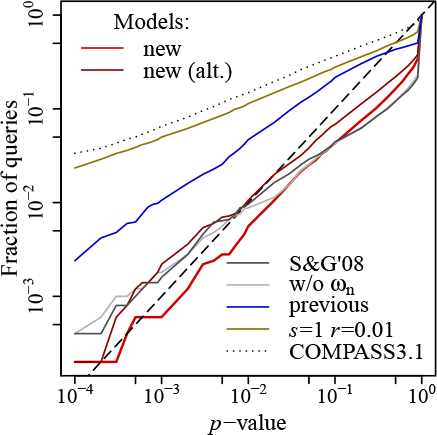
Statistical accuracy of the COMER method using different models for statistical significance estimation. Shown are the results of comparing the profiles constructed for 5000 randomized Pfam families to the profiles constructed for 4931 randomized SCOPe domains. The figure plots the fraction of queries with a *p*-value reported for their best match less than the *p*-value indicated on the *x*-axis. The dashed line indicates the highest accuracy. These models are represented in the figure: the new model (new) proposed, the alternative model with *E*-value correction (new (alt.)), a statistical model based on previous research (S&G’08), the statistical model used in the previous COMER version (previous), a model based on the statistics obtained for profiles generated with *s* = 1 and *r* = 0.01, and the model excluding the number of positive substitution scores (w/o *ω*_n_). The statistical accuracy of COMPASS (v3.1) is also shown.

The results are shown in Figure 4. The new model for statistical significance estimation is more accurate than the model (Margelevičius and Venclovas, 2010) implemented in the previous version (Margelevičius, 2018) of the COMER method. The results also show that the statistics obtained from aligning profiles generated by randomly permuting MSA columns (*s* = 1, *r* = 0.01) lead to overestimation of statistical significance. This result can be accounted for by unrealistic representation of protein sequence families (profiles).

Finally, although the models with and without the *ω*_n_ statistic taken into consideration achieve comparable statistical accuracy, the sensitivity and high-quality alignment rate obtained using the latter model (w/o *ω*_n_) are lower. The same applies to the model implemented based on the previous research (S&G’08) (see Section 3.2).

### 3.2 Sensitivity and alignment quality

We compared the performance of the new COMER version with that of its previous version (Margelevičius, 2018), where the new and previous versions differed only in how they estimated the statistical significance of the same alignments. For reference, we also provide results for three other profile-profile alignment methods, HHsearch (Söding, 2005) (v3.0.0), FFAS (Jaroszewski *et al.*, 2011), and COMPASS (v3.1) (Sadreyev and Grishin, 2008).

4900 protein domains of the test dataset from the SCOPe database (v2.03) filtered to 20% sequence identity was used to evaluate performance. The test and training datasets shared no common folds.

Profiles were constructed using two categories of MSAs. The MSAs for each domain sequence were obtained by running PSI-BLAST (Altschul *et al.*, 1997) (v2.2.28+) for six iterations and HHblits (Remmert *et al.*, 2012) for three iterations, respectively, against a filtered UniProt database (Suzek *et al.*, 2015).

A pair of aligned domains (profiles) that belonged to the same SCOPe superfamily or shared statistically significant structural similarity (DALI (Holm *et al.*, 2008) Z-score ≥ 2) was considered a true positive (TP). Aligned pairs that did not meet the above criteria but belonged to the same SCOPe fold were considered to have an unknown relationship and were ignored. Other aligned pairs were considered false positives (FPs). The sensitivity was also summarized using the ROC_*n*_ score, which is the normalized area under the ROC curve up to *n* FPs.

Alignment quality was evaluated by generating, using MODELLER (Šali and Blundell, 1993) (v9.4), protein structural models for each produced alignment. Statistically significant similarity between a model and the real structure, TM-score ≥ 0.4 (Zhang and Skolnick, 2004), was considered to correspond to a high-quality alignment (HQA). An alignment with a TM-score < 0.2, a characteristic value for a random pair, was assumed to be of low-quality (LQA).

Two evaluation modes were used to evaluate alignment quality. The local mode penalizes alignment overextension, whereas the global mode penalizes too short alignments (e.g., a few amino acids in length). These evaluation modes were also used to evaluate the quality of maximally extended alignments produced by COMER and HHsearch (option -mact set to 0). The evaluation of maximally extended alignments reveals how the quality of local alignments changes with their extension. It provides an indication of the quality of the match between the query and a database protein, which is also important in protein homology modeling.

(Details about the evaluation setting can be found in Supplementary Section S5.3.)

Figure 5 and Table 1 reveal that the new statistical model leads to consistent improvement in both sensitivity and HQA rate and that most of the improvements are statistically significant.

Using the developed statistical model yielded an increase of up to 34.2% and 27.4% in the number of TPs and up to 43.9% and 61.8% in the number of HQAs.

**Table 1.**
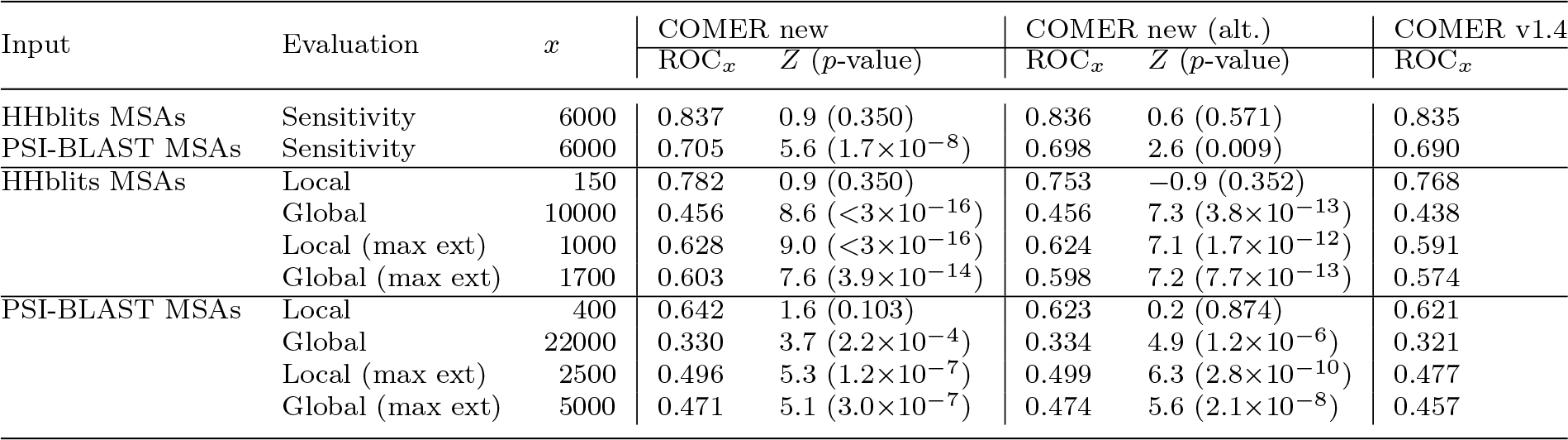
Area under the ROC curve and improvement of the COMER method. The sensitivity and alignment quality (Local and Global evaluation modes) of versions of the COMER method are evaluated. Maximally extended (max ext) COMER alignments are included in the evaluation. Profiles were constructed from HHblits and PSI-BLAST MSAs. ROC_*x*_ is the ROC score calculated up to *x* false positives (FPs; Sensitivity) or low-quality alignments (LQAs; alignment quality in the Local and Global modes). The number of FPs or LQAs, *x*, depends on the evaluation mode (see also Figure 5). *Z* is the difference between the areas (ROC_*x*_ scores) obtained for a new and the previous (v1.4) versions of the COMER method, divided by the estimated standard error. The statistical significance of *Z* is indicated in parentheses.

| Input | Evaluation | $x$ | COMER new | | COMER new (alt.) | | COMER v1.4 |
| --- | --- | --- | --- | --- | --- | --- | --- |
| | | | ROC<sub>$x$</sub> | $Z$ ($p$-value) | ROC<sub>$x$</sub> | $Z$ ($p$-value) | ROC<sub>$x$</sub> |
| HHblits MSAs | Sensitivity | 6000 | 0.837 | 0.9 (0.350) | 0.836 | 0.6 (0.571) | 0.835 |
| PSI-BLAST MSAs | Sensitivity | 6000 | 0.705 | 5.6 ($1.7 \times 10^{-8}$) | 0.698 | 2.6 (0.009) | 0.690 |
| HHblits MSAs | Local | 150 | 0.782 | 0.9 (0.350) | 0.753 | -0.9 (0.352) | 0.768 |
| | Global | 10000 | 0.456 | 8.6 ($<3 \times 10^{-16}$) | 0.456 | 7.3 ($3.8 \times 10^{-13}$) | 0.438 |
| | Local (max ext) | 1000 | 0.628 | 9.0 ($<3 \times 10^{-16}$) | 0.624 | 7.1 ($1.7 \times 10^{-12}$) | 0.591 |
| | Global (max ext) | 1700 | 0.603 | 7.6 ($3.9 \times 10^{-14}$) | 0.598 | 7.2 ($7.7 \times 10^{-13}$) | 0.574 |
| PSI-BLAST MSAs | Local | 400 | 0.642 | 1.6 (0.103) | 0.623 | 0.2 (0.874) | 0.621 |
| | Global | 22000 | 0.330 | 3.7 ($2.2 \times 10^{-4}$) | 0.334 | 4.9 ($1.2 \times 10^{-6}$) | 0.321 |
| | Local (max ext) | 2500 | 0.496 | 5.3 ($1.2 \times 10^{-7}$) | 0.499 | 6.3 ($2.8 \times 10^{-10}$) | 0.477 |
| | Global (max ext) | 5000 | 0.471 | 5.1 ($3.0 \times 10^{-7}$) | 0.474 | 5.6 ($2.1 \times 10^{-8}$) | 0.457 |

**Figure 5.**
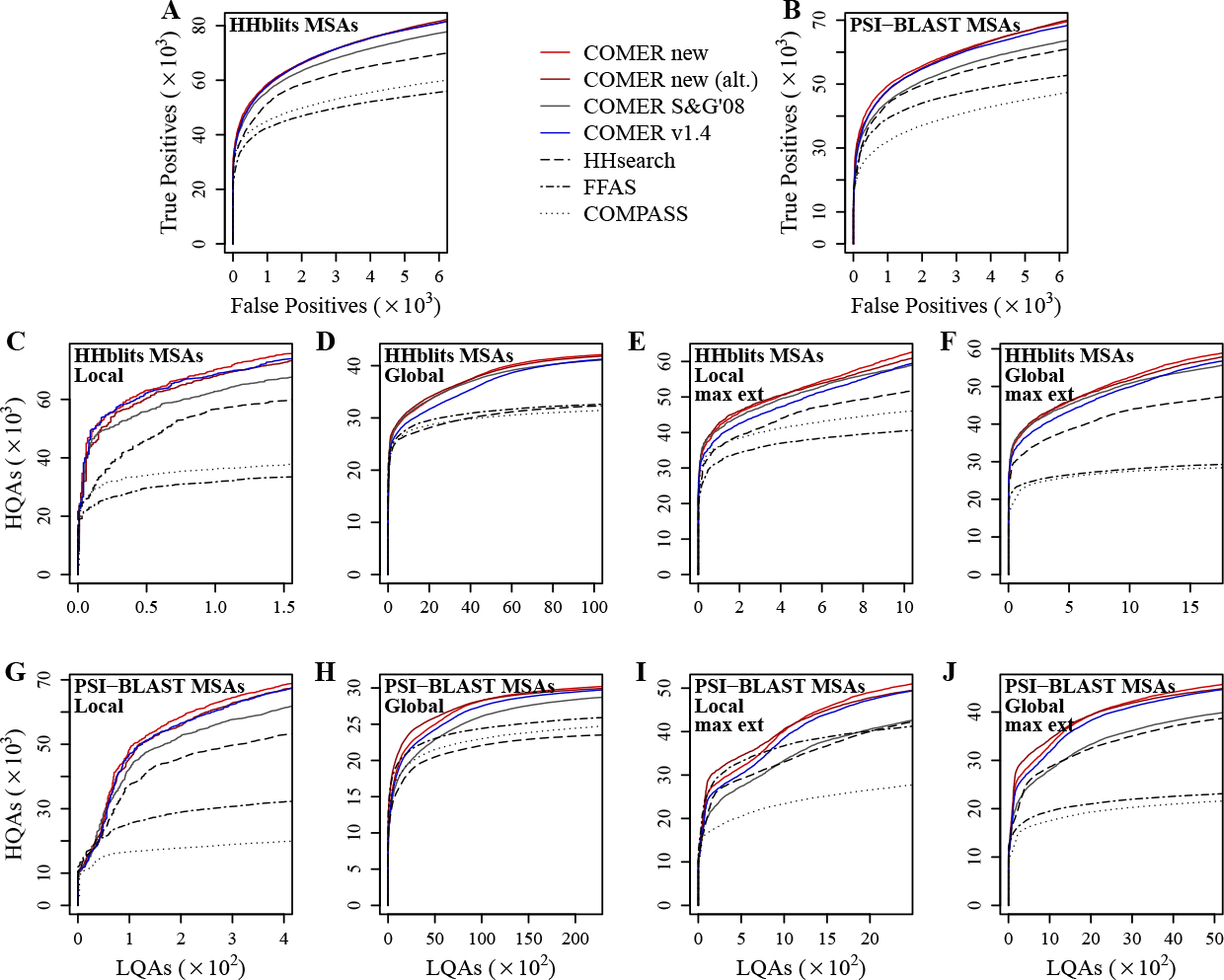
Performance of profile-profile alignment methods. (A,B): Sensitivity. (C–J): Alignment quality evaluated in the (C,E,G,I) local and (D,F,H,J) global evaluation modes. Maximally extended (max ext) COMER and HHsearch alignments are evaluated in (E,F,I,J). Profiles were constructed from (A,C–F) HHblits and (B,G–J) PSI-BLAST MSAs. COMER new (alt.) represents the COMER method using the model with *E*-value correction. COMER S&G’08 implements a statistical model based on previous research. COMER v1.4 represents the previous version of the COMER method. The color coding matches that of Figure 4. HQA, high-quality alignment. LQA, low-quality alignment.

Table 2 shows that these percentage increases were achieved at a low false discovery rate (FDR). Hence, the largest relative improvements are expected for relationships detected at a high confidence level.

Table 2 also shows that the relative increases were greater when the profiles were constructed from the PSI-BLAST MSAs. A similar trend was also observed for the number of HQAs examined as a function of the number of alignments of inferior quality (IQAs) (Supplementary Section S7.1). In this evaluation setting, the developed statistical model showed an increase of up to 102.8% and 193.3% in the number of HQAs.

MSAs obtained from a PSI-BLAST search contain more alignment errors than those built using HHblits. Larger relative improvements in performance achieved when using PSI-BLAST MSAs for profile construction, therefore, show a certain degree of robustness that the new model exhibit. We attribute this characteristic to the combination of statistical parameters dependent upon both profile length and ENO and compositional similarity. If a high alignment score of two unrelated profiles arises due to false positives in the corresponding MSAs, a high compositional similarity between the profiles counterbalances the statistical significance estimated solely based on profile length and ENO.

By contrast, the version based on the previous approach for estimating statistical significance (S&G’08) displays a large decrease in sensitivity and HQA rate with respect to the previous COMER version (v1.4) for PSI-BLAST MSAs. Among other differences from the new model, that model does not depend explicitly on the compositional similarity between profiles, which in part accounts for this decrease. Although version S&G’08 achieved similar statistical accuracy (Figure 4), the sensitivity and the rate of HQAs were consistently lower than those obtained using the new statistical model. (We also provide statistical analysis with respect to FPs found among the top-ranked alignments of the queries from the test dataset, but these results should be interpreted with caution [see Supplementary Section S7.2].)

### 3.3 Application to pairwise profile HMM alignments

We applied the methodology for estimating statistical significance (Figure 3) to pairwise profile HMM alignments produced by HHsearch. An improvement in both statistical accuracy and sensitivity and HQA rate (Supplementary Section S7.3) confirms the effectiveness of the developed methodology.

### 3.4 Example of reduced significance for a false positive

Reranking alignments using the proposed estimation of statistical significance has been shown to increase sensitivity and the rate of HQAs. This is demonstrated by an example.

Two domains d2hi7b1 (a.29.15.1) and d2cfqa (f.38.1.2) are alpha-helical proteins but represent different SCOPe classes. They have different topology and do not share statistically significant structural similarity.

Alpha-helical structure implies a high compositional similarity *λ*_u_ = 0.296 between the profiles constructed for the two domains (low values of *λ*_u_ correspond to high compositional similarity; Supplementary Sections S6.1 and S6.2). Significance estimation dependent upon compositional similarity allowed the new COMER version to correctly remove the alignment from the list of statistically significant alignments. In contrast, the alignment was considered significant by the previous COMER version. Note that both versions estimated the significance of the same alignment with the same score.

**Table 2.**
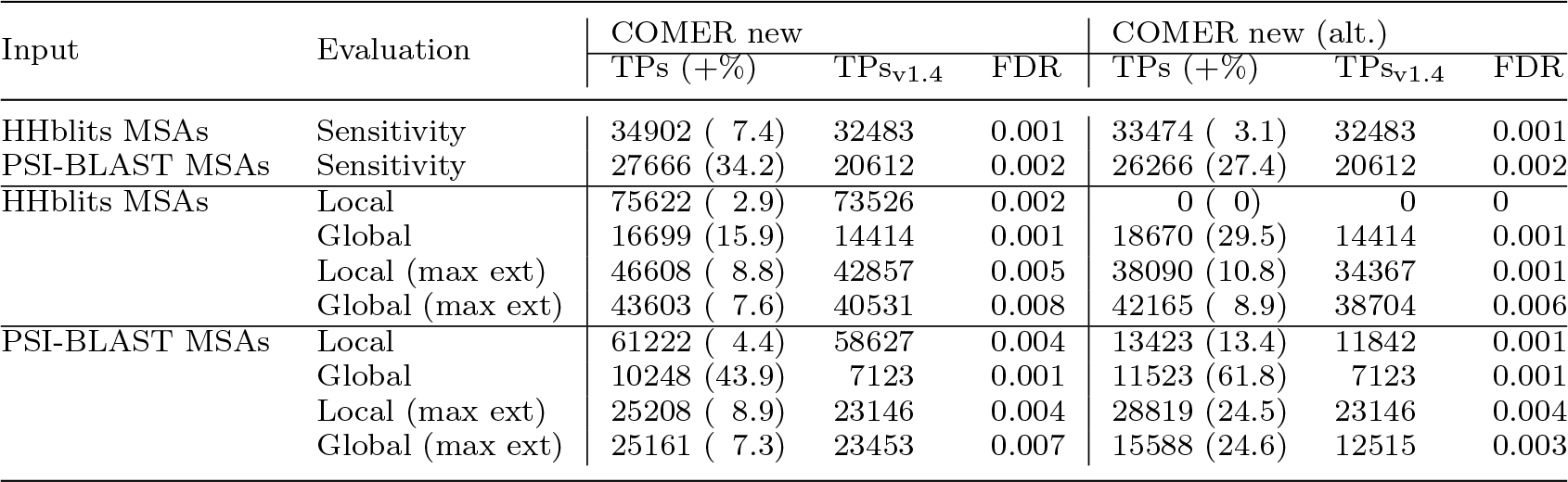
Increase in the number of true positives or high-quality alignments. Shown are the results of the evaluation of the sensitivity and alignment quality (Local and Global evaluation modes) of versions of the COMER method using profiles constructed from HHblits and PSI-BLAST MSAs. TPs stands for the number of true positives (Sensitivity) or high-quality alignments (in the Local and Global evaluation modes) at a specified false discovery rate (FDR) for a new version of the COMER method. TPs_v1.4_ represents the same number for the previous COMER version. The percentage improvement with respect to TPs_v1.4_ is given in parentheses. 0 indicates no improvement.

## 4 Discussion

Profiles represent sequence families, and this fact alone suggests that a profile contains information whose content largely depends on the family the profile describes. It means that the degree of (dis)similarity (distance) between unrelated sequence families strongly affects the distribution of scores of alignments between profiles that represent these families. The challenging aspects of characterizing the distribution of profile-profile alignment scores, however, are not limited to diversity across unrelated profiles. Similarity between SS, context and other predictions that accompany profiles complicate the characterization of alignment score distribution too.

The importance of a null model of profiles motivated us to develop an algorithm for generating random profiles. According to the algorithm, profiles are generated by concatenating fixed-length fragments sampled randomly from real unrelated profiles (seeds) with added noise. Controlling the noise level and fragment length allows to produce random profiles that are neither overly divergent nor share an excessive similarity. In this way, random profiles possess features characteristic to real profiles, but correlations that do not allow them to be considered unrelated do not dominate. Alignments between randomly generated profiles allowed us to determine the dependence of statistical parameters on profile length and ENO and also on compositional similarity between profiles.

The results suggest two important implications. First, profiles generated using long fragments (fragment length 9) represent real unrelated profiles much more accurately than do those obtained by randomly sampling profile columns. Statistics obtained from their alignments lead to higher statistical accuracy. On the other hand, randomly sampling of columns destroys higher-order dependencies inherent in real proteins and leads to the opposite result.

Second, the compositional similarity between profiles has a large impact on estimating the statistical significance of alignments. Improvements in sensitivity and HQA rate can be accounted for in part by that a high compositional similarity may indicate that the proteins share common structural elements or the profiles involve false positives. Significance estimation dependent upon compositional similarity, therefore, has a positive effect.

In fact, the issues of profile composition and random profile model are factors that hinder the application of techniques such as importance sampling (Park *et al.*, 2009) to accurately estimating statistical parameters. While composition can be measured in different ways, choosing a random profile model is more complicated because every profile represents a model. For example, the composition of two profiles, one of them obtained by randomly rearranging the positions of the other, will be the same. However, the distribution of alignment scores of profiles generated using the profile with rearranged positions as a seed (model) will differ from that obtained for profiles generated using the other profile as a seed. In this study, close match to the distribution of alignment scores of real unrelated reference profiles constituted one of the criteria for random profile model selection. However, a supplementary approach might be beneficial.

Based on the above considerations and the results obtained by generating random profiles using one seed profile, we suspect that further improvements may result from introducing a new measure of profile “sequence affinity” (as opposed to the quantitative measure of the effective number of observations). The benefit may come from including into the model the statistical parameters dependent upon the new measure through, e.g., the application of the conditional mean estimator. The present study already demonstrated the conditional mean estimator to be both easily interpretable and effective when the calculation of compositional similarity or divergence between profiles was in use. And it provides possibilities for further improvements.

## Funding

This research was funded by the European Regional Development Fund according to a supported activity under Measure No. 01.2.2-LMT-K-718 (Grant No. 01.2.2LMT-K-718-01-0028).

## References

Altschul, S. and Gish, W. (1996). Local alignment statistics. Methods Enzymol, 266, 460–480.

Altschul, S., Madden, T., Schäffer, A., Zhang, J., Zhang, Z., Miller, W., et al. (1997). Gapped BLAST and PSI-BLAST: a new generation of protein database search programs. Nucleic Acids Res, 25(17), 3389–3402.

Altschul, S., Bundschuh, R., Olsen, R., and Hwa, T. (2001). The estimation of statistical parameters for local alignment score distributions. Nucleic Acids Res, 29(2), 351–361.

Arratia, R. and Waterman, M. (1994). A phase transition for the score in matching random sequences allowing deletions. Ann Appl Probab, 4(1), 200–225.

Chernobai, A., Rachev, S., and Fabozzi, F. (2015). Composite goodness-of-fit tests for left-truncated loss samples. In C. Lee and J. Lee, editors, Handbook of Financial Econometrics and Statistics, pages 575–596. Springer, New York.

Dembo, A., Karlin, S., and Zeitouni, O. (1994). Limit distribution of maximal non-aligned two-sequence segmental score. Ann Probab, 22(4), 2022–2039.

Eddy, S. (2008). A probabilistic model of local sequence alignment that simplifies statistical significance estimation. PLoS Comput Biol, 4(5), e1000069.

Finn, R., Coggill, P., Eberhardt, R., Eddy, S., Mistry, J., Mitchell, A., et al. (2016). The Pfam protein families database: towards a more sustainable future. Nucleic Acids Res, 44(D1), D279–D285.

Fox, N., Brenner, S., and Chandonia, J. (2013). SCOPe: Structural classification of proteins–extended, integrating SCOP and ASTRAL data and classification of new structures. Nucleic Acids Res, 42(D1), D304–D309.

Holm, L., Kääriäinen, S., Rosenström, P., and Schenkel, A. (2008). Searching protein structure databases with DaliLite v.3. Bioinformatics, 24(23), 2780–2781.

Jaroszewski, L., Li, Z., Cai, X. H., Weber, C., and Godzik, A. (2011). FFAS server: novel features and applications. Nucleic Acids Res, 39, W38–W44.

Karlin, S. (2005). Statistical signals in bioinformatics. Proc Natl Acad Sci USA, 102(38), 13355–13362.

Karlin, S. and Altschul, S. (1990). Methods for assessing the statistical significance of molecular sequence features by using general scoring schemes. Proc Natl Acad Sci USA, 87(6), 2264–2268.

Karlin, S. and Brendel, V. (1992). Chance and statistical significance in protein and dna sequence analysis. Science, 257(5066), 39–49.

Karlin, S. and Dembo, A. (1992). Limit distributions of maximal segmental score among markov-dependent partial sums. Adv Appl Probab, 24(1), 113–140.

Karlin, S., Dembo, A., and Kawabata, T. (1990). Statistical composition of high-scoring segments from molecular sequences. Ann Stat, 18(2), 571–581.

Kotz, S. and Nadarajah, S. (2000). Extreme value distributions: theory and applications. Imperial College Press, London.

Margelevičius, M. (2016). Bayesian nonparametrics in protein remote homology search. Bioinformatics, 32(18), 2744–2752.

Margelevičius, M. (2018). A low-complexity add-on score for protein remote homology search with COMER. Bioinformatics, 34(12), 2037–2045.

Margelevičius, M. and Venclovas, Č. (2010). Detection of distant evolutionary relationships between protein families using theory of sequence profile-profile comparison. BMC Bioinformatics, 11, 89.

Meng, L., Sun, F., Zhang, X., and Waterman, M. (2011). Sequence alignment as hypothesis testing. J Comput Biol, 18(5), 677–691.

Messer, P., Bundschuh, R., Vingron, M., and Arndt, P. (2007). Effects of long-range correlations in DNA on sequence alignment score statistics. J Comput Biol, 14(5), 655–668.

Metzler, D. (2006). Robust E-values for gapped local alignments. J Comput Biol, 13(4), 882–896.

Mott, R. (1992). Maximum-likelihood estimation of the statistical distribution of smith-waterman local sequence similarity scores. Bull Math Biol, 54(1), 59–75.

Mott, R. (2000). Accurate formula for P-values of gapped local sequence and profile alignments. J Mol Biol, 300(3), 649–659.

Park, Y., Sheetlin, S., and Spouge, J. (2009). Estimating the Gumbel scale parameter for local alignment of random sequences by importance sampling with stopping times. Ann Stat, 37(6A), 3697–3714.

Pearson, W. (1998). Empirical statistical estimates for sequence similarity searches. J Mol Biol, 276(1), 71–84.

Poleksic, A. (2009). Island method for estimating the statistical significance of profile-profile alignment scores. BMC Bioinformatics, 10, 112.

Poole, W., Gibbs, D., Shmulevich, I., Bernard, B., and Knijnenburg, T. (2016). Combining dependent p-values with an empirical adaptation of Brown’s method. Bioinformatics, 32(17), i430–i436.

Remmert, M., Biegert, A., Hauser, A., and Söding, J. (2012). HHblits: lightning-fast iterative protein sequence searching by HMM-HMM alignment. Nat Methods, 9(2), 173–175.

Sadreyev, R. and Grishin, N. (2003). Compass: a tool for comparison of multiple protein alignments with assessment of statistical significance. J Mol Biol, 326(1), 317–336.

Sadreyev, R. and Grishin, N. (2008). Accurate statistical model of comparison between multiple sequence alignments. Nucleic Acids Res, 36(7), 2240–2248.

Schäffer, A., Aravind, L., Madden, T., Shavirin, S., Spouge, J., Wolf, Y., et al. (2001). Improving the accuracy of PSI-BLAST protein database searches with composition-based statistics and other refinements. Nucleic Acids Res, 29(14), 2994–3005.

Söding, J. (2005). Protein homology detection by HMM-HMM comparison. Bioinformatics, 21(7), 951–960.

Spang, R. and Vingron, M. (1998). Statistics of large-scale sequence searching. Bioinformatics, 14(3), 279–284.

Spang, R. and Vingron, M. (2001). Limits of homology detection by pairwise sequence comparison. Bioinformatics, 17(4), 338–342.

Suzek, B., Wang, Y., Huang, H., McGarvey, P., Wu, C., and the UniProt Consortium (2015). UniRef clusters: a comprehensive and scalable alternative for improving sequence similarity searches. Bioinformatics, 31(6), 926–932.

Šali, A. and Blundell, T. L. (1993). Comparative protein modelling by satisfaction of spatial restraints. J Mol Biol, 234(3), 779–815.

Waterman, M. and Vingron, M. (1994a). Rapid and accurate estimates of statistical significance for sequence data base searches. Proc Natl Acad Sci USA, 91(11), 4625–4628.

Waterman, M. and Vingron, M. (1994b). Sequence comparison significance and poisson approximation. Stat Sci, 9(3), 367–381.

Yu, Y. and Hwa, T. (2001). Statistical significance of probabilistic sequence alignment and related local hidden markov models. J Comput Biol, 8(3), 249–282.

Yu, Y., Gertz, E., Agarwala, R., Schäffer, A., and Altschul, S. (2006). Retrieval accuracy, statistical significance and compositional similarity in protein sequence database searches. Nucleic Acids Res, 34(20), 5966–5973.

Zhang, Y. and Skolnick, J. (2004). Scoring function for automated assessment of protein structure template quality. Proteins, 57(4), 702–710.

